# Diurnal dynamics of multilayer brain networks predict cognitive trajectories in aging

**DOI:** 10.1101/2025.03.24.644492

**Authors:** Kenza Bennis, Anna Canal-Garcia, Joana B. Pereira, Giovanni Volpe, Francis Eustache, Christophe Phillips, Christine Bastin, Fabienne Collette, Gilles Vandewalle, Thomas Hinault

**Affiliations:** Inserm, U1077, EPHE, UNICAEN, Normandie Université, PSL Université Paris, CHU de Caen, GIP Cyceron, Neuropsychologie et Imagerie de la Mémoire Humaine (NIMH), 14000 Caen, France; Department of Clinical Neuroscience, Karolinska Institutet, Stockholm, Sweden; Department of Physics, University of Gothenburg, Gothenburg, Sweden; GIGA-CRC-Human Imaging, Université de Liège and Belgian National Fund for Scientific Research, Liège, Belgium

**Keywords:** multilayer, integration, recruitment, rsFC, diurnal rhythms, cognitive aging, brain oscillations, circadian

## Abstract

**Objectives:** Resting-state functional connectivity (rsFC) is a highly dynamic process that varies across different times of the day within each individual. Although this variability was long considered to be noise, recent evidence suggests it may allow for an optimal adaptation to changes in the environment. However, the way rsFC is shaped on a circadian scale and its association with cognition are still unclear.

**Methods:** We analyzed data from 90 late-middle-age participants from the Cognitive Fitness in Aging study (61 women; 50-69y). Participants completed five electroencephalographic (EEG) recordings of spontaneous resting-state activity spread over 20h of prolonged wakefulness. Using a temporal multilayer network approach, we characterized the diurnal variations of the dynamic recruitment and integration of resting-state brain networks. We focused on the theta and gamma frequency bands within the default mode network (DMN), central executive network (CEN), and salience network (SN). Additionally, we investigated the relationship recruitment and integration of these network with baseline cognitive performance and at 7-year longitudinal follow-up, as well as with positron emission tomography (PET) early neuropathological markers of Alzheimer’s disease such as β-amyloid and tau/neuroinflammation.

**Results:** Diurnal changes in theta and gamma dynamics were associated with distinct cognitive aspects. Specifically, higher baseline memory performance was associated with higher theta dynamic integration of the SN and the CEN, as well as higher theta dynamic recruitment of the DMN. Moreover, lower longitudinal memory decline at 7-year was associated with higher theta dynamic integration of the SN, CEN, and DMN. In contrast, higher gamma diurnal dynamic integration of the SN and the CEN was associated with lower executive and attentional performance, as well as higher early β-amyloid accumulation, at baseline.

**Discussion:** These findings suggest that maintaining a balance between network flexibility and stability throughout the diurnal phase of the circadian cycle may play a crucial role in cognitive aging, with stable theta-band connectivity supporting memory, whereas excessive gamma-band stability in the SN and CEN may contribute to executive decline and early amyloid accumulation. These insights highlight the importance of considering time-of-day in brain rsFC studies, calling for a temporal multilayer approach to capture these dynamic patterns more effectively.

## 1. INTRODUCTION

One of the major challenges of cognitive neuroscience is to understand the functional organization of the brain underlying human cognitive abilities and behaviors. Beyond brain activations, brain networks are key to the organization of cognitive abilities (Miraglia et al., 2017, Chan et al., 2017). A functional brain network can be defined as a system composed of brain regions that are more functionally connected to each other within the system than to the other regions of the brain, a property referred to as network recruitment. At the same time, brain networks are part of a highly integrated large-scale network where between-network interactions, referred to as network integration, play a key role in orchestrating complex cognitive processes. Resting-state functional connectivity (rsFC) analyses of brain signal is considered to reflect the stable and intrinsic organization of brain networks that can be related to cognition (Yeo et al., 2011; Uddin et al., 2019). Most of these studies have analyzed rsFC in isolation and averaged over entire recording sessions (or scan), implicitly assuming that brain networks organization remains stable over space and time (Bassett et al., 2011, Calhoun et al., 2014). However, accumulating evidence suggests that brain networks continuously fluctuate, requiring a more dynamic perspective.

In fact, rsFC is inherently dynamic, fluctuating both over short and long timescales, ranging from moment-to-moment switching of brain regions’ connectivity patterns among various functional brain networks, to slow age-related reorganization. Together, these dynamic phenomena have proven promising in finding determinants of cognitive trajectories in aging (Sala-Llonch et al., 2015). At the level of a single scan, short-term fluctuations in rsFC, particularly the frequency with which brain regions switch between different networks, have been shown to predict cognitive performance (Pedersen et al., 2018). At the longest timescale, across the lifespan, rsFC undergoes gradual reorganization, reflecting compensatory mechanisms that sustain cognitive function despite age-related non-lesional changes. In particular, it has been shown that a hallmark of aging is increased integration and decreased recruitment of brain networks’ rsFC associated with preserved cognitive performance. This indicates a shift toward a more distributed network organization and reduced functional specialization of brain networks to maintain cognitive performance (Varangis et al., 2019; He et al., 2020). However, because these slow changes accumulate over years, they cannot directly capture the ongoing neural dynamics underlying cognitive variability (Jockwitz & Caspers, 2021). Between these two extremes, an intermediate timescale exists: the diurnal phase of the circadian cycle.

Here, we investigate age-related brain changes associated with diurnal changes in brain networks dynamics at rest. Unlike aging-related changes, diurnal fluctuations occur within hours and can be directly measured, offering a unique opportunity to examine the temporal organization of rsFC associated with the large heterogeneity of individual cognitive trajectories observed during aging (Anderson et al., 2017, Cabeza et al., 2018). The rsFC signals follow structured variations across the day, shaped by the interplay between prior sleep-wake history, which sets the need for sleep, and the circadian system, which promotes wakefulness and cognition during the day while favoring sleep at night (Gaggioni et al., 2019, Ly et al., 2016). However, the diurnal rsFC fluctuations in cognitive aging remains largely unexplored. In a first step toward understanding diurnal rsFC fluctuations, we previously used electroencephalography (EEG) to investigate how rsFC varies across five different times of the day (five EEG sessions), and how these fluctuations relate to cognition and tau and β-amyloid biomarkers in healthy late middle-aged participants (Bennis et al., 2024). EEG time-frequency analyses, with its millisecond-level temporal resolution, is uniquely suited to capture subtle changes in spontaneous neural dynamics over time and detect early signs of age-related cognitive decline (Courtney & Hinault, 2021). For each session, we measured rsFC as the synchronization of brain rhythm between brain regions, with each rhythm associated with different cognitive processes and defined by its corresponding frequency band, from the slowest to the fastest rhythm: delta (1-4Hz), theta (4-8Hz), alpha (8-12Hz), beta (13-30Hz) and gamma (30-100Hz). To quantify diurnal rsFC fluctuations, we computed a global fluctuation index across all five sessions. Among the key findings, we observed that greater diurnal rsFC fluctuations in theta band was associated with poorer memory performance and higher tau/neuroinflammation levels, whereas greater diurnal rsFC fluctuations in gamma band correlated with better executive function and lower β-amyloid deposition. Couplings involved in these associations were part of the salience network (SN) and the central executive network (CEN). While this previous work highlighted that diurnal rsFC fluctuations in the theta and gamma bands may reflect critical processes underlying cognitive aging, it did not capture the specific mechanisms of network reorganization, nor did it consider longitudinal cognitive trajectories.

Here, to address these gaps, we applied multilayer network analyses, a method well-suited for capturing rsFC dynamics over time (Mucha et al., 2010). In this approach, each layer represents a time-specific rsFC pattern, forming a time-varying functional network that models the continuous reorganization of brain connectivity (Sporns, 2018). Unlike classical methods, this framework allows us to differentiate dynamic recruitment (the stability of functional associations within a network), from dynamic integration (the extent to which regions switch between networks over time) (Mattar et al., 2015). We investigated how these dynamics relate to 6-to-7-year longitudinal cognitive trajectories and tau/β-amyloid biomarkers in healthy late middle-aged adults. Specifically, we examined SN, CEN, and DMN recruitment and integration across the day, as these networks play a central role in cognition and aging-related changes (Menon, 2011; La Corte et al., 2016). We expected increased integration relative to recruitment, indicating reduced functional specialization throughout the day.

We further explored how diurnal patterns of recruitment and integration relate to cognitive trajectories and pathological markers. In the theta band, we hypothesized that greater dynamic integration would be linked to better cognitive performance, slower cognitive decline, and lower tau/neuroinflammation, reflecting preserved brain efficiency. In contrast, in the gamma band, we expected that greater dynamic integration would be associated with poorer cognitive performance, greater decline, and higher β-amyloid levels, suggesting a maladaptive process. Given that participants showed no marked cognitive deterioration, these associations may reflect early compensatory mechanisms in healthy aging preceding pathological deposition.

## 2. RESULTS

We analyzed data from 90 healthy late middle-aged adults (50–69 years old) from the COFITAGE database (COgnitive FITness in AGEing) database [see Materials and Methods and (Van Egroo et al., 2019, 2021; Narbutas et al., 2021; Chylinski et al., 2021, 2022; Rizzolo et al., 2021; Bennis et al., 2024)], who underwent five resting-state EEG recordings over the course of a single day (10 a.m., 4 p.m., 8 p.m., 10 p.m., and 1 a.m.). rsFC was measured using the Phase Lag Index (PLI), which quantifies synchronization between brain regions, in theta (4–8 Hz) and gamma (30–100 Hz) bands. To capture rsFC dynamics in theta and gamma bands, we constructed two time-varying multilayer networks, where each PLI matrix represents a layer reflecting the rsFC pattern at a given time, allowing us to measure dynamic recruitment and integration. Higher rsFC fluctuations corresponded to lower recruitment and integration scores, indicating reduced network stability. In addition to EEG, cognitive assessments at baseline and after 7 years, along with tau and β-amyloid markers, were collected to examine the associations between rsFC, cognitive trajectories, and pathological aging processes.

### 2.1. Networks’ diurnal recruitment and integration dynamics are similar in theta and gamma

We performed a repeated measures ANOVA (Networks x Dynamic Measures x Frequencies) to determine how the regions of our 3 brain networks of interest were recruited and integrated over the day in both theta and gamma frequency bands. The analysis first yielded a significant effect of dynamic measure (F (1,170) =347.489, p<0.001, ηp² = 0.803), with higher dynamic recruitment compared to integration of brain networks’ regions over time (Figure 1). The interaction between network and dynamic measure (F (2,170) =245.431, p<0.001, ηp² =0.743), revealed that the recruitment of the SN (M=0.464, SE=0.006) is significantly higher than the recruitment of the CEN (M=0.450, SE=0.005), which is also significantly higher than the recruitment of the DMN (M=0.415, SE=0.004) (Figure 1.B). Moreover, dynamic integration was significantly higher for DMN brain regions (M=0.405, SE=0.004) than for both the SN (M=0.402, SE=0.004) and the CEN brain regions (M=0.399, SE=0.004) (Figure 1.A). No interaction with the frequency band was found, indicating that these patterns of results did not significantly differ between theta and gamma bands.

**Figure 1.**
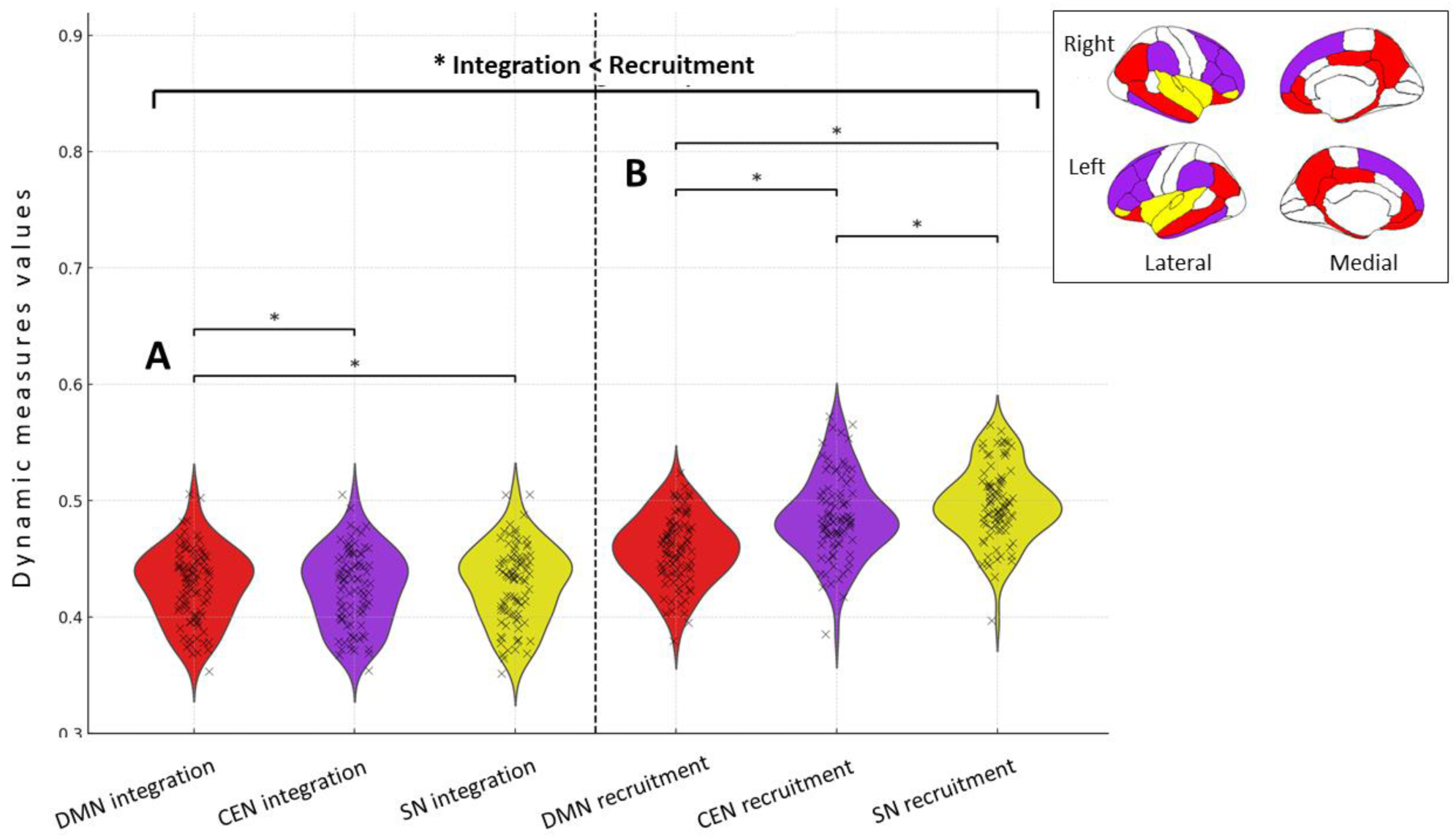
Dynamic recruitment is higher than integration across DMN, SN, and CEN in theta and gamma bands. Violin plots illustrate higher dynamic recruitment (right) than integration (left) across DMN (red), SN (yellow), and CEN (purple) over time in the theta and gamma bands. **A)** Integration: DMN (M=0.405, SE=0.004) was significantly higher than both SN (M=0.402, SE=0.004) and CEN (M=0.399, SE=0.004). **B)** Recruitment: SN (M=0.464, SE=0.006) was significantly higher than CEN (M=0.450, SE=0.005), which was in turn higher than DMN (M=0.415, SE=0.004). B) Integration: DMN (M=0.405, SE=0.004) was significantly higher than both SN (M=0.402, SE=0.004) and CEN (M=0.399, SE=0.004). Brain maps (top right) highlight the corresponding regions in each network.

### 2.2. Theta and gamma diurnal dynamic network measures are differently associated with cognitive performance

To test whether diurnal dynamic measures (integration and recruitment) could explain part of individual differences in cognitive performance among older individuals, we assessed their associations with cognitive performance at baseline and with cognitive decline after 7-year follow-up. Specifically, we assessed how dynamic recruitment and integration of the SN, CEN and DMN in both theta and gamma bands related to three domain-specific composite scores, the recognition memory score (MST), and their corresponding cognitive decline scores. For all the results below, no effect of age, sex and mean gray matter volume or individuals’ wake-up/DLMO phase value was found, indicating that these factors did not significantly contribute to the observed associations. The correlation matrix in Figure 2.C provides Pearson’s R-values of the associations assessed through regression analyses.

**Figure 2.**
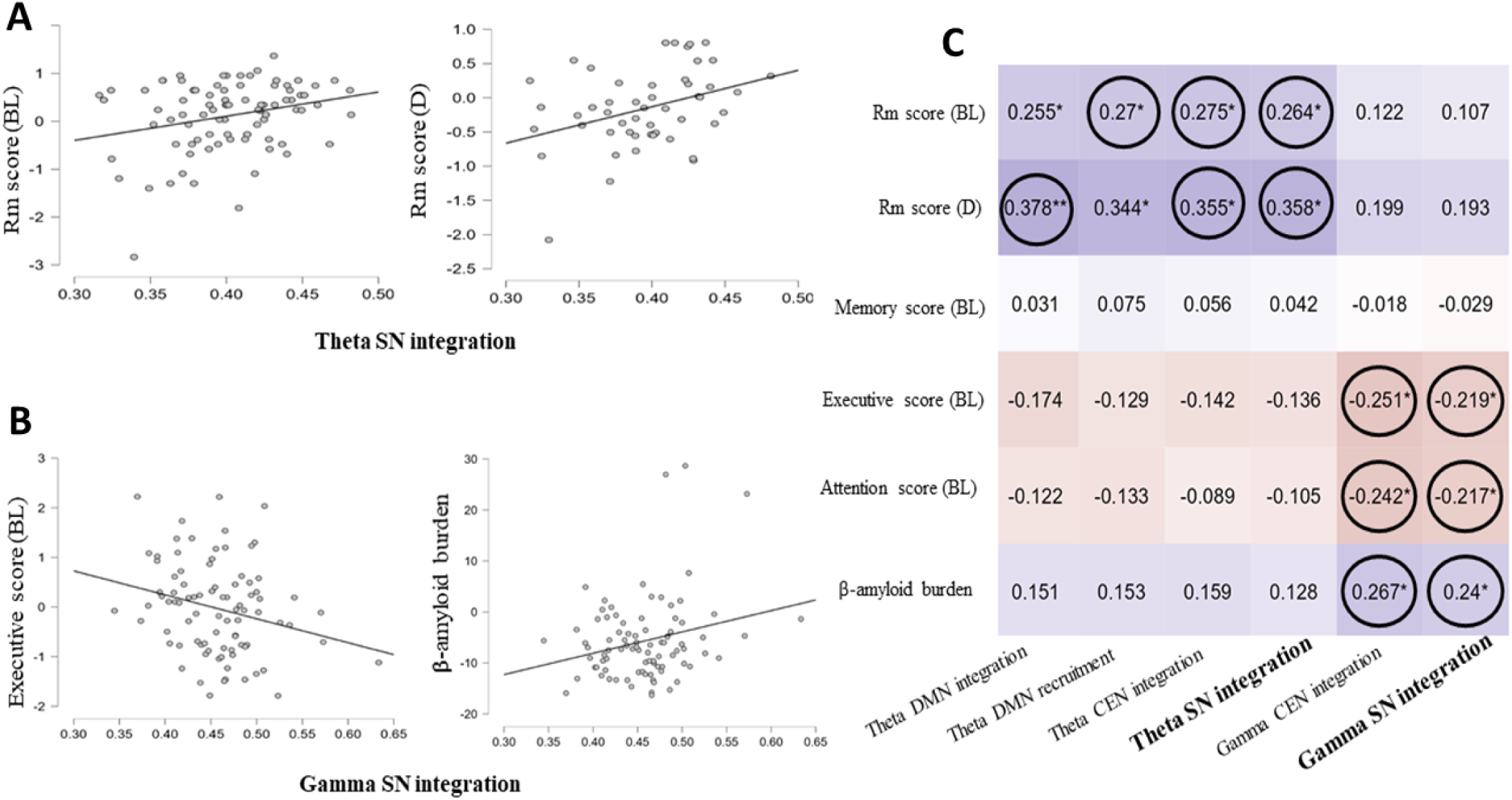
Distinct associations between theta and gamma SN integration, cognitive performance, and β-Amyloid burden. Scatter plots illustrate key associations between SN integration and cognitive performance. **A)** In theta band, higher SN integration is associated with higher memory performance at baseline (left) and with lower longitudinal memory decline (right). **B)** In constrast, in the gamma band, higher SN integration is associated with lower executive performance at baseline (left) and higher β-amyloid burden (right). **C)** Correlation matrix shows R-values of Pearson’s correlations (*: p<.05, **: p<.01; positive correlation: blue, negative correlation: red), black circles indicate results that survived when FDR corrected for multiple comparisons (Benjamini & Hochberg, 1995). BL = baseline cognitive performance; D = decline in performance between baseline and 7 years follow-up.

#### 2.2.1. Positive association of diurnal dynamic DMN recruitment and diurnal dynamic SN and CEN integration in theta band with memory performance at baseline

Regression analyses showed that DMN diurnal dynamic recruitment and SN and CEN diurnal dynamic integration coefficients in theta band were positively correlated with the recognition memory score of the MST at baseline (DMN: β = 5.506, t = 2.580, p=0.012, FDR-corrected p = 0.0325; SN: β = 5.049, t = 2.505, p=0.014, FDR-corrected p = 0.04; CEN: β = 5.442, t = 2.621, p=0.010, FDR-corrected p = 0.04). These results indicate that higher dynamic recruitment of DMN and higher dynamic integration of SN and CEN brain regions across the day in theta band are associated with higher memory performance at baseline (Figure 2.A left).

#### 2.2.2. Negative association of SN and CEN diurnal dynamic integration in gamma band with executive and attention performance at baseline

Regression analyses showed that SN and CEN diurnal dynamic integration coefficients in gamma band were negatively correlated with the executive composite score (SN: β = -4.818, t = -2.376, p=0.020, FDR-corrected p = 0.04; CEN: β = -5.275, t = -2.678, p=0.009, FDR-corrected p = 0.018) and the attention composite score (SN: β = -5.627, t = -2.609, p=0.011, FDR-corrected p = 0.04; CEN: β = -5.993, t = -2.858, p=0.005, FDR-corrected p = 0.018) both at baseline. In other words, higher dynamic integration of SN and CEN brain regions across the day in gamma band are associated with lower executive and attention performance at baseline (Figure 2.B left).

#### 2.2.3. Positive association of diurnal dynamic DMN, SN and CEN integration in theta band with longitudinal memory decline

Regression analyses showed that DMN, SN and CEN diurnal dynamic integration coefficients in the theta band, were positively correlated with the recognition memory score of the MST decline score (DMN: β = 5.682, t = 2.825, p=0.007, p=0.011, FDR-corrected p = 0.028; SN: β = 5.328, t = 2.657, p=0.011, FDR-corrected p = 0.044; CEN: β = 5.334, t = 2.633, p=0.011, FDR-corrected p = 0.012). That is, higher dynamic integration of DMN, SN and CEN brain regions across the day in the theta band are associated with a lower memory decline after 7 years follow-up (Figure 2.A right).

#### 2.2.4. Positive association between SN and CEN diurnal dynamic integration in gamma band and early β-amyloid burden

Results showed that SN and CEN diurnal dynamic integration coefficients in the gamma band were positively correlated with β-amyloid burden rate (SN: β = 51.179, t = 2.493, p=0.015, FDR-corrected p = 0.023; CEN: β = 54.585, t = 2.744, p=0.007, FDR-corrected p = 0.009). Higher dynamic integration of SN and CEN brain regions across the day in the gamma band is associated with higher β-amyloid burden (Figure 2.B right).

## 3. DISCUSSION

Brain region’s rsFC variability is increasingly considered to be predictive of cognitive performance in aging (Garrett et al., 2011, Courtney & Hinault, 2021; Jauny et al., 2022; Hinault et al., 2021, 2023; Uddin et al., 2020; Kumral et al., 2020), reflecting the dynamic reorganization of brain regions. Furthermore, the switching rates between different network communities have been associated with cognitive performance (Pedersen et al., 2018). Our main goal was therefore to characterize brain network’s rsFC diurnal dynamics in healthy cognitive aging. Using a multilayer network approach, we investigated theta and gamma rsFC variability across five recordings over 20h of continuous (mainly daytime) waking, in late middle-aged healthy participants (50-69y). Our results shed new light on a) how brain regions are integrated and recruited across the day in relation to their resting-state network and network communities, b) the associations of so-called diurnal dynamic networks’ recruitment and integration with cognitive performance, and c) their associations with cognitive decline and with biological markers of Alzheimer’s disease (AD).

### 3.1. Theta and gamma diurnal dynamic reorganization of DMN, SN and CEN

We first showed that DMN, SN, and CEN brain regions were more often recruited with regions belonging to the same network community than other communities across the day, both in theta and gamma band. In other words, over the day, rsFC of brain regions tended to remain within the same community rather than to switch between communities. This finding suggests that functional specialization of brain networks is preserved over the course of the day, while the integration between different networks fluctuates to a greater extent. This interpretation appears to contrast with the findings of a previous fMRI study that compared network dynamics between morning and evening sessions in young adults (Farahani et al., 2022). Their results showed greater integration than recruitment in the evening compared to the morning, which was interpreted as a reflection of functional dedifferentiation occurring over the day. However, their conclusions were based on a comparison of rsFC at only two discrete time points and did not capture the continuous evolution of functional network organization across the day. Our study provides additional information about extended diurnal dynamics of rsFC brain networks as our multilayer model accounts for five different times of the day ordered in a 20-hour timeframe. By using a finer-grained temporal resolution through the incorporating of intermediate recordings between morning and evening, we reveal that brain networks maintain a stable recruitment pattern throughout the day, preserving their functional specialization, while integration fluctuates more dynamically. These additional time points offer new insights into the temporal evolution of rsFC and highlight the importance of considering fine-grained temporal resolution when studying brain network dynamics.

Diurnal dynamic recruitment of brain regions over time indicated that rsFC reorganization, or rsFC variability, over the day was higher within the SN than the CEN and then the DMN. Previous studies showed that the SN is involved in the regulation of arousal, as rsFC alterations within the SN were reported in individuals with sleep disturbance (Ma et al., 2020, Khazaie et al., 2017). Thus, high rsFC reorganization within the SN might reflect the progressive build-up of sleep need during prolonged wakefulness. Conversely, lower rsFC within the DMN during the day has been associated with lower maintenance of wakefulness and might also reflect the build-up of sleep pressure (Facer-Childs et al., 2019). Lower reorganization of DMN brain regions rsFC might indicate its lower implication as the day progresses. Brain regions of the CEN exhibit an intermediate rate of rsFC reorganization across the day, in line with previous studies characterizing the CEN as the most stable resting-state network across the day (Blautzik et al., 2013). Finally, our results showed that diurnal dynamic integration of brain regions over time was higher for the DMN than for SN and CEN, indicating that over the day, DMN regions exhibited greater reorganization with other network communities compared to SN and CEN regions. This result is consistent with previous studies in healthy young adults, showing that higher integration of DMN regions with other networks over the day is associated with deliberate mind-wandering and introspective processes (Golchert et al., 2017, Farahani et al., 2022). The DMN, SN, and CEN play a crucial role in cognitive aging due to their involvement in key cognitive processes such as attention, memory, and executive functions (Menon et al., 2011, 2023). The SN regulates the balance between DMN activity, which supports internally directed cognition, and CEN activity, which is involved in goal-directed tasks such as working memory and attentional control (Chen et al., 2013). Aging has been associated with changes in the functional organization of these networks, including increased integration among them (Ng et al., 2016), which has been interpreted as a compensatory mechanism to counteract cognitive decline. Specifically, studies have shown that altered SN-DMN interactions are observed in individuals with mild cognitive impairment (Chand et al., 2017), and that reorganization of these large-scale networks is a key marker of cognitive aging (La Corte et al., 2016). Taken together, our results suggest that the diurnal dynamic recruitment and integration of the DMN, SN, and CEN, networks that are critical for maintaining cognitive function in aging.

### 3.2. High diurnal theta SN, CEN and DMN dynamics are associated with better memory performance at baseline and lower longitudinal memory decline

Our results reveal that higher diurnal dynamic integration of both SN and CEN brain regions, and higher diurnal dynamic recruitment of DMN brain regions, all in theta band, were associated with higher memory performance at baseline. This concerns the recognition memory score of the MST, which has been shown to be a sensitive measure to detect subtle general cognitive decline in aging (Chylinski et al. 2022, Rizzolo et al., 2021, Pishdadian et al., 2020).

Results concerning SN and CEN are in line with previous studies, showing increased integration of brain networks during aging, associated with similar cognitive performance as younger adults (Stanley et al., 2015; Droby et al., 2022; Jockwitz & Caspers, 2021). These positive associations with cognitive performance could reflect a compensatory response to reduced network specialization in older adults (Damoiseaux et al., 2017; Wig et al., 2017; Setton et al., 2023; Malagurski et al., 2020; Koen et al., 2019; He et al., 2020). However, our results might indicate that this compensation is most effective when brain network integration remains stable throughout the day. The increased integration of SN and CEN may therefore reflect an adaptive reorganization of large-scale network dynamics, helping to counteract the decline in network segregation commonly observed in aging (Damoiseaux et al., 2017; Wig et al., 2017; Setton et al., 2023; Malagurski et al., 2020; Koen et al., 2019; He et al., 2020). In this context, our findings suggest that, rather than transient increases in rsFC, maintaining stable integration of SN and CEN regions in theta band throughout the day is essential for sustaining memory performance. Unlike SN and CEN regions, DMN regions appears to maintain stable within-network rsFC throughout the day in theta band, which is also associated with preserved memory performance at baseline. This finding is in line with a study showing that, while SN and CEN exhibit early functional changes and increased inter-network connectivity in normal cognitive aging, the DMN appears to maintain more sustained within-network functional connectivity (Oschmann et al., 2020). As the largest brain network, the DMN consists of key hub regions involved in numerous cognitive processes, including memory (Ferreira et al., 2013). Aging induces important reorganization of the rsFC between these hub regions to compensate the loss of efficiency of these regions to maintain cognitive performance (Andrews Hanna et al., 2010, Ferreira et al., 2013). Altogether, our results suggest that, in healthy cognitive aging, diurnal dynamic integration of SN and CEN regions might reflect a beneficial ability of brain network to reorganize across the day with other networks to maintain cognitive performance, whereas DMN brain regions remain functionally consistent across the day.

Furthermore, higher diurnal dynamic integration of SN, CEN and DMN regions, in theta band, was associated with lower memory decline after a 7-year follow-up. These findings refine our understanding of the adaptive mechanisms that support cognitive preservation in aging, by emphasizing the diurnal temporal dynamics of this compensatory process. Rather than a simple increase in inter-network connectivity, the ability to maintain stable rsFC patterns across the day appears to be a critical determinant of cognitive trajectories in aging. Specifically, stable inter-network rsFC throughout the day, particularly in the theta band, appears to be beneficial for memory in older adults, whereas fluctuations in inter-network connectivity may indicate a failure to sustain alternative compensatory networks, as it is associated with long-term memory decline. These findings suggest that cognitive benefits depend on efficient network communication, not just broader engagement (Cabeza, 2018; Reuter-Lorenz & Park, 2014). A stable rsFC throughout the day, particularly in the theta band, may thus reflect optimal network efficiency and be associated with better memory preservation in aging.

Finally, our results highlight the central role of theta-band rsFC dynamics in memory and cognitive aging. Previous research has shown that aging is associated with an increase in slow rhythms activity, including theta, which reflects a general slowing of brain activity and has been linked to cognitive decline (Lopez et al., 2014; Jensen et al., 2014). Notably, theta-band rsFC has been identified as a key predictor of cognitive decline, including the progression from subjective cognitive complaints to mild cognitive impairment (MCI), and from MCI to Alzheimer’s disease (Babiloni et al., 2021). Theta-band have long been associated with memory processes and large-scale network coordination (Fell & Axmacher, 2011), and enhanced theta rsFC has been linked to preserved memory performance in aging (Finnigan & Robertson, 2011). While no significant differences were observed across frequency bands in our study, the associations between diurnal rsFC dynamics and memory performance were exclusively linked to theta-band phase synchrony. This suggests that, within the theta frequency range, diurnal dynamic integration is associated with healthy cognitive trajectories in aging, regardless of the specific network involved.

### 3.3. High diurnal gamma SN and CEN dynamic integration are associated with low executive and attentional performance at baseline

In contrast, higher diurnal dynamic integration of SN and CEN regions in gamma band, reflecting greater stability of rsFC across the day, was associated with lower executive and attentional performance. Conversely, greater fluctuations in gamma rsFC were associated to better executive and attentional functioning, suggesting that flexibility, rather than stability, in gamma band supports cognitive efficiency. These findings align with previous studies indicating that excessive integration of SN and CEN in aging, when accompanied by reduced performance, may reflect maladaptive network reorganization rather than a compensatory mechanism (Ng et al., 2016). Moreover, a recent study reported that while localized gamma synchrony facilitates cognitive function, widespread gamma synchrony—akin to excessive connectivity stability—is associated with lower cognitive performance, particularly in tasks requiring high cognitive effort (Bakhtiari et al., 2023). This pattern is consistent with the neural dedifferentiation hypothesis which suggests that aging leads to a loss of functional specialization (Malagurski et al., 2020). However, a rigid gamma connectivity pattern appears detrimental to cognition. According to the Scaffolding Theory of Aging and Cognition (STAC; Reuter-Lorenz & Park, 2014), cognitive aging benefits from the adaptive engagement of alternative networks, highlighting the importance of gamma-band flexibility throughout the day.

This interpretation is supported by research showing that gamma oscillations play a key role in attentional control, cognitive flexibility, and information integration (Fries et al., 2005; Missonnier et al., 2010; Lundqvist et al., 2018). In aging, a high degree of flexibility in gamma connectivity may reflect an ability to dynamically allocate cognitive resources, while excessive stability in gamma synchrony may indicate reduced adaptability and impaired executive efficiency. However, unlike the theta band, gamma dynamics were not associated with long-term cognitive trajectories. While gamma fluctuations may enhance momentary executive function, they do not necessarily predict preserved cognition over time. This absence of a long-term effect raises the possibility that gamma connectivity reflects a state of hyperexcitability that could be neurotoxic in the long run (Drinkenburg, Tok & Ahnaou, 2022). While some level of gamma flexibility appears beneficial, excessive fluctuations or persistent instability could disrupt efficient network coordination, potentially contributing to cognitive decline.

### 3.4. High diurnal gamma SN and CEN dynamic integration are associated with high early β-amyloid burden rate

In the gamma band only, we observed that higher diurnal dynamic integration of SN and CEN regions was associated with higher early β-amyloid burden. This result can be linked to recent findings showing increased widespread gamma synchrony in AD patients (Basar et al., 2016), reinforcing the idea that rsFC stability in gamma band may be linked to early neuropathological processes. Considerable evidence suggests amyloid deposition precedes a decline in cognition and may be the initiator of a cascade of events that indirectly leads to cognitive decline (Rodrigue et al., 2009). Our results further refine this perspective by revealing that participants with greater rsFC stability in the gamma band throughout the day exhibited higher levels of early β-amyloid deposition, whereas those with greater fluctuations in gamma connectivity had lower β-amyloid burden. This finding suggests that a lack of flexibility in gamma connectivity may be an early biomarker of pathological changes in the brain. This is consistent with previous research indicating that the presence of β-amyloid proteins in their early deposition sites might disrupt the functional connectivity between large-scale networks, in particular the inter-connectivity of SN with other regions before the spread of the pathology that occurs later (Guzman-Vélèz et al., 2022).

This aligns with neurostimulation studies showing that increased gamma-band flexibility can promote amyloid clearance (Murdock et al., 2024). Future research should clarify whether rigid gamma connectivity serves as an early indicator of β-amyloid pathology, either as a marker of emerging deposition or as a downstream effect of amyloid-related disruptions. While β-amyloid accumulation has been predominantly linked to the DMN (Raichle, 2015), our findings highlight the role of diurnal SN–CEN integration in the gamma band, suggesting that amyloid-related processes may extend beyond the DMN.

### 3.5. Limitations and perspectives

Our findings have some limitations. First, our results were obtained from healthy middle-aged to older participants whose global cognitive performance is in the upper average range, and none of them exhibited significant cognitive decline during the follow-up period. Therefore, the present results could differ in individuals aged 70 and above or in those showing early signs of cognitive decline. Furthermore, the study did not include a group of young adults, so we were not able to assess group differences relative to younger participants to determine the age-specificity of our results. However, our study included a longitudinal aspect, which accounts for individuals’ cognitive performance seven years after baseline and imply that some the links we report are related to within subject modification. Finally, two-minutes of EEG resting-state recording might not be deemed enough to optimally characterize the ongoing brain state. Recent studies showed however that resting-state brain activity allows the robust differentiation of individuals from brain recordings as short as 30 seconds, ushering the notion of neural fingerprint (Castanheira and al., 2021).

## 4. CONCLUSION

rsFC variability, long considered as noise, has been identified as a key marker of age-related cognitive variability in several BOLD signal MRI studies (Garrett et al., 2011), whereas its characterization through EEG remains underexplored, limiting our understanding of its electrophysiological underpinnings in aging. Here, we reported a diurnal pattern of dynamic SN, CEN, DMN rsFC associated with baseline cognitive performance, longitudinal cognitive trajectories and pathological markers, which differ when considering theta or gamma band, suggesting that these two rhythms have specific and opposite roles in cognitive functioning and AD pathology. In the theta band, greater stability in network integration was associated with better memory performance, suggesting that a well-maintained connectivity structure supports efficient long-range communication. In contrast, in the gamma band, greater fluctuations in network integration were linked to better executive and attentional function, indicating that these cognitive abilities benefits from more flexible connectivity patterns. These findings highlight that neither stability nor flexibility is inherently beneficial, their effects depend on the underlying frequency band and cognitive function involved. More broadly, our findings suggest that diurnal rsFC dynamics may parallel aging-related changes in brain function, offering a compressed timescale representation of the aging process. The association between cognitive performance and diurnal network dynamics parallels the long-term increase in network integration observed in aging, often interpreted as an adaptive response to declining neural efficiency (Malagurski et al., 2020). Investigating rsFC dynamics within the CEN, DMN, and SN on a diurnal scale, as described in the triple network model (Menon, 2011), provide new insights into cognitive aging.

Taken together, our findings support the promises of diurnal rsFC variability as a marker of cognitive functioning in aging. They also highlight the value of temporal multilayer network approaches using EEG time-frequency data in capturing subtle brain long before cognitive changes in aging. Future research should refine these findings and explore their clinical applications, particularly for identifying early markers of neurodegeneration.

## ACKNOWLEDGEMENTS

This research was not preregistered and did not receive any specific grant from funding agencies in the public, commercial, or not-for profit sectors. Data collection was conducted at the GIGA-In Vivo Imaging platform of ULiège, Belgium. Data sharing for this project were provided by the GIGA-CRC Human Imaging. Data access can be provided upon reasonable request to FC and GV.

Funding for this project was provided by Fonds National de la Recherche Scientifique (FRS-FNRS, FRSM 3.4516.11, F.4513.17 and T.0242.19, EOS Project MEMODYN No. 30446199; Belgium), the Wallonia-Brussels Federation (Grant for Concerted Research Actions— SLEEPDEM 17/27–09), Stop Alzheimer foundation (Belgium, grants 15018, 2019/0025), University of Liège, Fondation Simone et Pierre Clerdent, European Regional Development Fund (ERDF, Radiomed Project). [18F]Flutemetamol doses were provided and cost covered by GE Healthcare Ltd (Little Chalfont, UK) as part of an investigator sponsored study (ISS290) agreement. This agreement had no influence on the protocol and results of the study reported here.

JBP and ACG were supported by Swedish Research Council (#2022-01108), the Swedish Alzheimer Foundation (#AF-968323), a Consolidator Karolinska Institute grant, the Swedish Brain Foundation (FO2022-0147), Gamla Tjänarinnor (#2020-01016; #2021-01207; #2022-01341), KI foundations, Stohnes or the project “A Multimodal Brain Connectivity Marker for the Early Detection of Alzheimer’s Disease” funded by European Union – NextGenerationEU and the Romanian Government, under National Recovery and Resilience Plan for Romania, contract no. 760250/28.12.2023, cod PNRR-C9-I8-CF109/31.07.2023, through the Romanian Ministry of Research, Innovation and Digitalization, within Component 9, Investment I8.

## 5. METHODS

### 5.1. Participants

Participants were enrolled in a multimodal longitudinal study designed to identify cerebral biomarkers of normal cognitive aging (the Cognitive Fitness in Aging – COFITAGE – study; Van Egroo et al., 2019). We analyzed data from ninety healthy late middle-aged participants (61 women and 29 men; aged 50-69 years), with EEG data recorded for all five sessions. Based on previous work investigating aging effects on oscillatory activity, a sample size of ninety participants would provide 90% power to detect an effect size of Cohen’s f = 0.14 (medium effect size), with an alpha = 0.05 (Hinault et al., 2021). No participants reported any recent history of neurological or psychiatric disease or were taking medication affecting the central nervous system. All participants had anatomically normal MRI scans, with no significant grey or white matter abnormalities. Extended information about protocol, exclusion criteria, recruitment, consent and financial reward can be found in previous publications from this cohort (first in Van Egroo et al., 2019, 2021; Narbutas et al., 2021; Chylinski et al., 2021, 2022; Rizzolo et al., 2021, Bennis et al., 2024). A subsample of 64 participants who had data for tau/neuroinflammation-PET imaging was also considered for additional analyses. After a 7-years follow-up, 58 participants were seen again for a similar neuropsychological assessment. Demographic characteristics of the final samples at baseline and after the 7-year follow-up are described in Table 1. The study was approved by the Medicine Faculty/Hospital Ethics Committee of the University of Liège, Belgium.

**Table 1.** Demographic, cognitive, neuroimaging, and biomarker characteristics at baseline and 7-year follow-up. Demographic information, cognitive scores, tau (THK-PET in Braak I/II ROIs) and β-amyloid burden (in early-stage deposition) and structural measures are presented for the entire sample at baseline and for 58 participants after 7 years follow-up. PET imaging (tau and β-amyloid) was only conducted at baseline; values at follow-up correspond to the subset of participants who underwent PET at baseline and were reassessed cognitively after 7 years. Mean (standard deviations) are provided.

|  | Baseline | 7 years follow-up |
| --- | --- | --- |
| **NEUROPSYCHOLOGICAL ASSESSMENT** |  |  |
| **NUMBER OF PARTICIPANTS** | 90 | 58 |
| **NUMBER OF FEMALES, N (%)** | 60 (66.7%) | 36 (62.1%) |
| **AGE, MEAN (S.D.)** | 59.2 (5.289) | 67.3 (5.448) |
| **YEARS OF EDUCATION, MEAN (S.D.)** | 15.2 (3.009) |  |
| **MEMORY** |  |  |
| **COMPOSITE SCORE, MEAN (S.D.)** | 0.014 (0.925) | 0.108 (1.411) |
| FCSRT (SUM OF ALL FREE RECALLS), MEAN (S.D.) | 34.228 (5.159) | 33.017 (5.969) |
| MST (RECOGNITION MEMORY SCORE), MEAN (S.D.) | 0.792 (0.153) | 0.786 (0.183) |
| **EXECUTIVE** |  |  |
| **COMPOSITE SCORE, MEAN (S.D.)** | -0.038 (0.908) | -0.113 (2.475) |
| 2-MIN VERBAL LITERAL FLUENCY, MEAN (S.D.) | 24.660 (7.076) | 26.996 (7.312) |
| 2-MIN VERBAL CATEGORIAL FLUENCY, MEAN (S.D.) | 33.820 (7.032) | 33.724 (7.013) |
| DIGIT SPAN (INVERSE ORDER), MEAN (S.D.) | 6.614 (2.182) | 6.810 (2.290) |
| TMT (RT FOR PART B), MEAN (S.D.) | 68.673 (20.331) | 64.776 (20.817) |
| N-BACK (3-BACK VARIANT, D-PRIME), MEAN (S.D.) | 0.673 (0.145) | -0.115 (1.090) |
| **ATTENTION** |  |  |
| **COMPOSITE SCORE, MEAN (S.D.)** | 0.039 (0.981) | -0.112 (2.301) |
| DSST (2-MIN SCORE), MEAN (S.D.) | 72.560 (12.629) | 72.155 (11.309) |
| TMT (RT FOR PART A), MEAN (S.D.) | 31.703 (8.671) | 29.000 (8.615) |
| N-BACK (1-BACK VARIANT, D-PRIME), MEAN (S.D.) | 0.976 (0.041) | -0.112 (1.308) |
| D2 (GZ-F SCORE), MEAN (S.D.) | 401.730<br>(72.233) | 412.448<br>(61.843) |
| **AB AND TAU/NEUROINFLAMMATION – PET IMAGING** |  |  |
| **TAU/NEUROINFLAMMATION BURDEN** |  |  |
| NUMBER OF PARTICIPANTS | 64 | 40 |
| NUMBER OF FEMALES, N (%) | 42 (65.6%) | 25 (62.5%) |
| THK-PET IN BRAAK I/II ROIS, MEAN (S.D.) | 2.289 (0.247) | 2.307(0.277) |
| **B-AMYLOID BURDEN** |  |  |
| NUMBER OF PARTICIPANTS | 90 | 57 |
| NUMBER OF FEMALES, N (%) | 68 (67.3%) | 36 (63.2%) |
| B-AMYLOID EARLY STAGE, MEAN (S.D.) | 0.855 (0.052) | -5.590(8.794) |
| **MRI** |  |  |
| CORTICAL GRAY MATTER VOLUME (MM<sup>3</sup>), MEAN (S.D.) | 458647.230<br>(39834.230) |  |

### 5.2. Wake-extension protocol

Five EEG recordings of spontaneous resting-state activity were performed over the course of 15 hours, between 10a.m. and 1a.m. in the context of a wake-extension protocol. Participants were required to follow a regular sleep-wake schedule (±30 minutes) for 1 week based on their preferred bed and wake-up times prior to the study, in order to ensure standardized baseline conditions and minimize the impact of sleep variability on cognitive measures. Compliance was verified using sleep diaries and wrist actigraphy (Actiwatch, Cambridge Neurotechnology). The day before, participants arrived at the laboratory 8 hours before their habitual bedtime and were kept in dim light (<5 lux) for 6.5 h preceding bedtime. Following the baseline night of sleep, the wake-extension protocol consisted of 20 hours of continuous wakefulness under strictly controlled constant routine conditions, (i.e., in-bed semi-recumbent position, dim light <5 lux, temperature ∼19°C, regular isocaloric food intake, no time-of-day information and sound-proofed rooms) to counteract the effect of external influences on endogenous circadian rhythms, assessed with salivary melatonin. The time of all procedures of the wake extension protocol was adjusted according to the clock time of the participants’ habitual sleep-wake schedule. At baseline, neuropsychological assessment, β-amyloid-PET and Tau/neuroinflammation-PET imaging together with T1-weighted MRI were also acquired on separate visits. Only neuropsychological assessment was re-performed after 7-years follow-up (Figure 3A).

**Figure 3.**
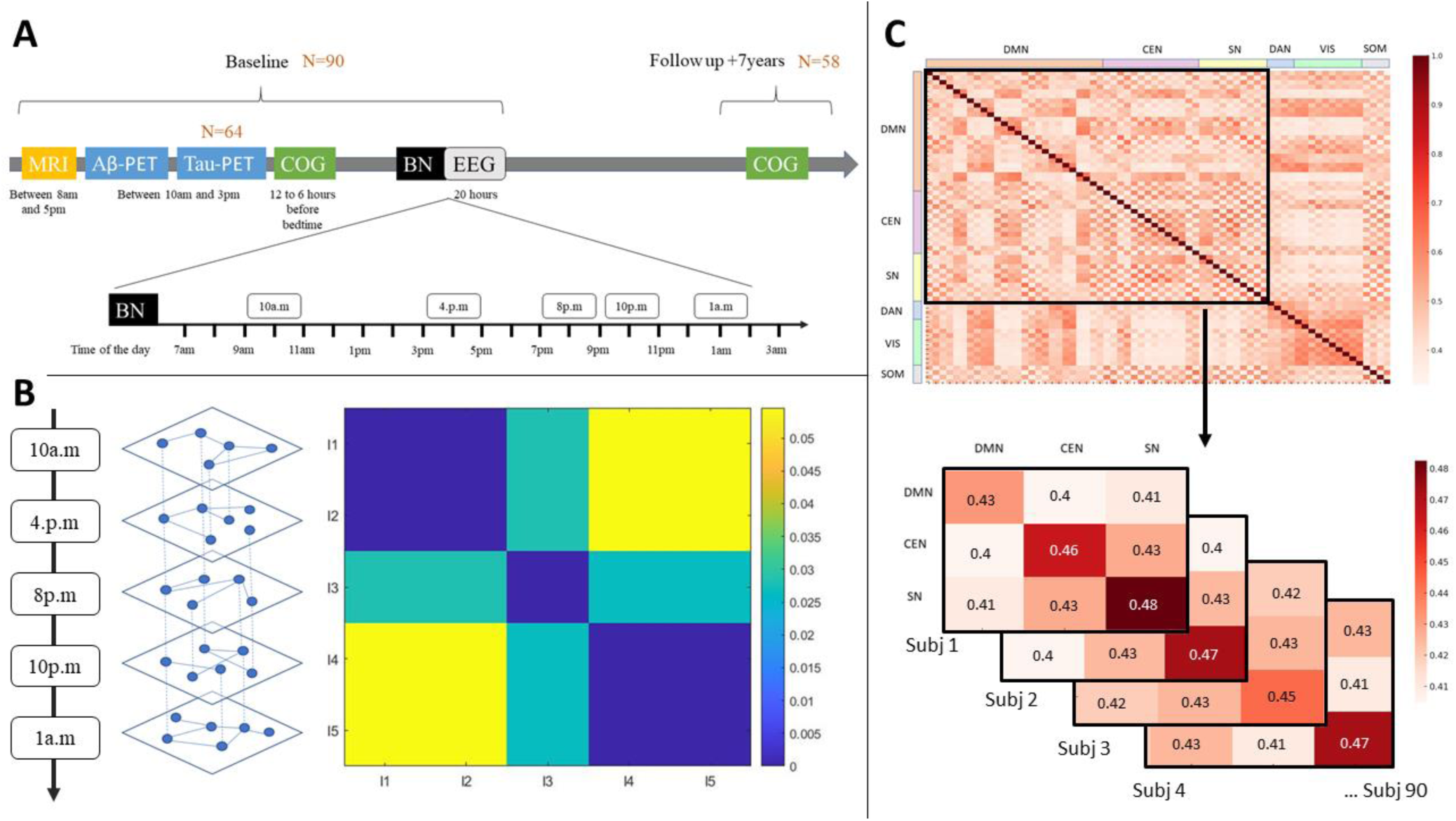
Experimental protocol and multilayer network analysis of diurnal rsFC dynamics. **A)** Experimental protocol, adapted from Van Egroo et al. (2019). COG: cognitive assessment, BN: Baseline-Night. **B)** Representation of the temporal multilayer network and Variation of Information (VI) matrix (ω = 0.5 and γ = 1.03) at the group level. For each participant, we constructed two temporal multilayer networks (theta and gamma), composed of five functional layers of PLI matrices, featuring the 5 EEG sessions in their chronological order. In each layer, nodes were connected by edges only to their corresponding nodes in other layers in a consecutive or chronological order. VI matrix showed how community structure change over layers at group-level, and revealed 3 time-periods in the daily organization of brain regions communities at group level. **C**) Module allegiance matrix of brain regions (top), averaged by networks (bottom), both for each participant. The diagonal of the matrices represents the dynamic recruitment and the off-diagonal represents the dynamic integration.

### 5.3. Neuropsychological assessment

At the first visit, neuropsychological assessment was administered in two 1.5h sessions and consisted of a battery of cognitive tests assessing three specific domains: memory, attention and executive functions. The raw scores were converted to z-scores and three domain-specific composite scores were computed as the standardized sum of z-scores of the domain-specific scores, where higher values indicate better performance. As part of the memory assessment, we also considered the recognition memory score of the Mnemonic Similarity Task (MST; Stark et al., 2013), which evaluates the ability to visually recognize images of objects that were incidentally encoded. Previous published work on the COFITAGE database (Chylinski et al. 2022, Rizzolo et al., 2021), in line with the literature (Pishdadian et al., 2020), showed that this MST score might be an early cognitive marker of memory decline. Analyses focused on the three composite scores and the MST recognition memory score. Seven-years later, a similar follow-up neuropsychological assessment was administered and the same procedure was used to compute the three composite scores. For each score, a cognitive decline score was calculated as the baseline performance minus the follow-up performance, divided by the baseline performance, so that a negative score reflects a decline over the 7 years and a positive score reflects a cognitive gain (all the cognitive scores are detailed in Table 1).

### 5.4. Salivary melatonin assessment

Salivary melatonin was measured by radioimmunoassay (Stockgrand Ltd, Guildford, UK). The detection limit of the assay for melatonin was 0.8 ± 0.2 pg/l using 500 µl volumes. To account for the fact that each participant’s circadian phase is different, and that the course of the protocol might vary slightly between individuals, we considered the Dim-light melatonin onset time (DLMO). DLMO were computed for each participant using the hockey-stick method, with ascending level set to 2.3 pg/ml (Hockey-Stick software v1.5). Saliva samples were collected every hour in order to specify individuals’ endogenous circadian rhythmicity during time awake by computing the phase between individuals’ wake-up time and individuals’ DLMO time (DLMO = phase 0°; 15° = 1h), since we were interested in fluctuations over the course of the day.

### 5.5. PET imaging

Amyloid-β PET and Tau/neuroinflammation-PET imaging were performed on an ECAT EXACT+ HR scanner (Siemens, Erlangen, Germany). β-Amyloid-PET imaging was performed with radiotracers [18F]Flutemetamol except for three subjects for which [18F]Florbetapir was used. Tau/neuroinflammation-PET imaging was performed with [18F]THK5351 for all subjects. For both β-Amyloid and tau/neuroinflammation PET imaging, a standardized uptake value ratio (SUVR) was calculated (Table 1). As β-Amyloid-PET imaging were acquired using different radioligands, their SUVR values were converted into Centiloid units (in line with previous works of this cohort, see Van Egroo et al., 2019 and Narbutas et al., 2021). Volumes of interest were determined using the automated anatomical labeling atlas (AAL; Tzourio-Mazoyer et al., 2002). β-Amyloid burden was averaged over composite masks covering neocortical regions reported to undergo the earliest aggregation sites for β-Amyloid pathology (frontal medial cortex, fusiform gyrus and temporal gyrus), while Tau/neuroinflammation burden was averaged over regions corresponding to Braak stages of early regional tau pathology (entorhinal cortex and hippocampus). The detailed PET imaging procedure was previously published in Narbutas et al., 2021.

### 5.6. Anatomical data

Participants’ T1-weighted MRI acquisition was performed on a 3Tesla MR scanner (MAGNETOM Prisma, Siemens) to assess brain grey matter integrity (Table 1). The following parameters were used: repetition time (TR) = 18.7ms; flip angle (FA) = 20 degrees; 3D multiecho fast low angle shot (FLASH) sequence (TR/FA) = 136166; voxel size = 1 mm3 isotropic; acquisition time of all structural sequences = 19 minutes (see Van Egroo and al., 2019 for detailed parameters). The FreeSurfer (Fischl, 2012) software was used to generate cortical surfaces and automatically segment cortical structures from each participant’s T1-weighted anatomical MRI, to account for individual brain anatomy during source reconstruction.

### 5.7. EEG recording and analyses

#### 5.7.1. Data acquisition

For each participant, two minutes of resting-state EEG (sampling rate: 1450 Hz, bandpass filter: 0.1-500 Hz) were recorded five times throughout the wake-extension protocol (for example at 10:00 a.m., 4:00 p.m., 8:00 p.m., 10:00 p.m. and 1:00 a.m. for a participant waking up at 7:00am) with a 60-channel EEG system (Eximia, Nexstim, Helsinki, Finland) covering the whole scalp. Participants were instructed to relax and avoid blinking while staring at a black dot.

#### 5.7.2. Pre-processing

Artifact and channel rejection (on continuous data), filtering (0.5-40Hz bandpass, on unepoched data), re-referencing (i.e., using the algebraic average of the left TP9 and right TP10 mastoid electrodes) and source estimation were performed using Brainstorm (Tadel et al., 2011). Physiological artefacts (blinks, saccades) were identified and manually removed through Independent Component Analyses (ICA) using Infomax algorithm (EEGLAB, runica.m). Independent Component Analyses approach consists in removing artifacts from the recording without removing the affected data portions, by identifying spatial components that are independent in time and uncorrelated with each other (Tadel et al., 2011). Source estimates were aligned to a standard space (i.e., ICBM152 template) using Brainstorm’s automated spatial registration pipeline with individual FreeSurfer surfaces. This ensured accurate cortical surface alignment across participants.

#### 5.7.3. Sources reconstruction

FreeSurfer (Fischl, 2012) segmentation of individuals T1-weighted anatomical MRI was used to constraint the source reconstruction and account for individual anatomy. The EEG forward model was obtained from a symmetric boundary element method (BEM model; OpenMEEG, Gramfort et al., 2010), fitted to the spatial positions of each electrode. A cortically constrained sLORETA procedure (Pascual-Marqui and Lehmann, 1994) was applied to estimate the cortical origin of scalp EEG signals. The estimated sources were then projected onto the ICBM152 standard space for comparisons between groups and individuals, while accounting for differences in native anatomy.

#### 5.7.4. PLI matrices construction

The individual alpha-peak frequency (IAF) observed at occipital sites was used to estimate the range of each frequency band. Based on previous works (Toppi et al., 2018; Hinault et al., 2021) the following five frequency bands were considered: Delta (IAF – [6–8] Hz), Theta (IAF – [2– 6] Hz), Alpha (IAF + [-2–2] Hz), Beta (IAF + [2–14] Hz), and Gamma1 (IAF + [15–30] Hz).

Phase-lag index (weighted PLI analyses; Stam et al., 2007) was used to assess the phase synchrony between 68 regions of interest (ROI; 68 ROIs as 34 contralateral homologous ROIs) defined by using the Desikan-Kiliany brain atlas (Desikan et al., 2006). For each participant, we obtained 5 x 5 PLI matrices of 68×68 ROIs couplings, with one matrix per frequency band for each of the five EEG recording sessions. PLI analyses estimate the variability of phase differences between two regions over time. Similar phase difference across time is indicated by a PLI value close to 1 (i.e., high synchrony between regions), while large variability in the phase difference is indicated by a PLI value close to 0. PLI measure has been shown to be less sensitive to the influence of common sources and amplitude effects relative to phase-locking value, as it disregards zero phase lag that could reflect volume conduction artefacts (Stam et al., 2007). Following analyses are focused only on the Theta and Gamma frequency band as our previous study showed their specific implication in diurnal brain dynamics in healthy aging (Bennis et al., 2024).

### 5.8. Multilayer network analyses

#### 5.8.1. Temporal multilayer network

Using BRAPH 2 software (Mijalkov et al., 2017; https://github.com/braph-software/BRAPH-2/releases/tag/2.0.0.a4) and following a procedure detailed in Canal-Garcia et al. (2024), we created two temporal multilayer networks for each participant, one for the theta and another one for the gamma frequency band. Each temporal multilayer network is composed of five “functional” layers of 68×68 PLI matrices, always featuring the five EEG sessions in their chronological order (Pipeline Ordered-Multiplex Connectivity Analysis using Weighted Undirected graphs). In each layer, nodes were connected by edges only to their corresponding nodes in other layers in a consecutive or chronological order (Figure 3B).

#### 5.8.2. Multilayer community structure

A community detection approach (Mucha et al., 2010) was used to cluster groups of nodes that are more highly connected to one another than to nodes outside their communities across the five layers, by computing the generalized multilayer modularity *Q* as follows: 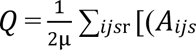 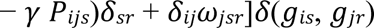, where µ is the total weights of the edges, *A*_*ijs*_ is the adjacency matrix between nodes *i* and *j* at layer *s*, γ is the resolution parameter, which sets the weights of intralayer connections at layer *s*, *P*_*ijs*_ is the associated null matrix (i.e. edges randomization of each layers while maintaining node strength) at layer *s*, δ_*sr*_ = 1 if *s* = *r* and 0 otherwise, ω is the temporal resolution parameter which determines the weights of the inter-layer edges, δ_*ij*_ = 1 if *i* = *j* and 0 otherwise, *g*_*is*_ and *g*_*jr*_ are the community allegiances of node *i* at layer *s* and node *j* and layer *r* respectively, and δ(*g*_*is*_, *g*_*jr*_) = 1 if the community allegiances *g*_*is*_ and *g*_*jr*_ of nodes *i* and *j* at layer *s* and *r* are the same and 0 otherwise. Low γ values produced fewer but larger communities, while high values resulted in more but smaller communities. Small ω values emphasized unique community structures per layer, while larger values highlighted shared community structures across layers, representing potential community structures that do not change over time (Puxeddu et al., 2020, Canal-Garcia et al., 2024). We chose intermediate values, such as ω = 0.5 and γ = 1.03, following previous work comparing ω and γ values (Matter et al., 2015; Puxeddu et al., 2020), so as to maximize the variability of the flexibility coefficient across brain regions. At the group level, we obtained 3 community structures across the five temporal layers, in both theta and gamma, which have been used to initialize individuals’ analyses. Finally, to enhance the robustness of the communities found in each layer, we performed 100 iterations of the multilayer modularity optimization algorithm for each participant, with each iteration initialized using the group-level community structure. This iterative approach ensured that the final community assignments were stable and not dependent on a single initialization. Variation of Information (VI) matrices averaged across all participants were plotted to visualize how the community structure evolve over layers (i.e., over time) at group-level (Figure 3B). Each cell in the matrix represents the VI value between the community structures of the corresponding layers. Low VI values indicate that community structures are similar over layers reflecting a stable functional connectivity over time, whereas high VI values indicate that community structures are different over layers suggesting variability of functional connectivity over time. VI matrix revealed 3 time-periods in the diurnal organization of brain regions communities: 1) stable organization of brain regions communities between the first and the second layers and between the fourth and the fifth layers, 2) major reorganization of brain regions communities between the first two layers and the last two layers, 3) moderate changes in the organization of brain regions communities between the third layer and the other layers suggesting that the third layer might be a transition period.

#### 5.8.3. Module allegiance matrix and dynamic network measures

To further investigate the temporal dynamics of brain network modularity, we computed and plotted the module allegiance matrix for each participant, which measures the probability that a pair of nodes (two brain regions) are assigned to the same community over time and repetitions (Mattar et al., 2015) (Figure 3C, top). The matrix M consists of 68 x 68 pair of nodes and can be written as: 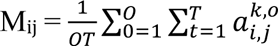, where O is the number of iterations of the multilayer community detection algorithm, T is the number of layers. For each optimization O and layers T, if they are in the same community network, the value of module allegiance is 1 (the values on the main diagonal of the matrix are all 1); otherwise, it is 0.

To quantify the dynamic role of a region within and between brain networks, we calculated dynamic network recruitment and the dynamic network integration coefficient based on the module allegiance matrix (Bassett et al., 2015; Mattar et al., 2015). For this purpose, six resting-state networks were predefined based on Uddin et al. (2019) classification: Central Executive Network (CEN), Salience Network (SN), Default Mode Network (DMN), Dorsal Attention Network (DAN), Visual System (VS) and Sensorimotor Network (SMN). Brain regions were assigned to these networks according to this classification, and module allegiance was subsequently calculated as a result of the community detection algorithm. Dynamic recruitment coefficient, was defined as the probability that a region is assigned to the same network community as other regions from the same network over layers and repetitions. For node *i* in the community network *S*, the dynamic recruitment can be computing as follows: 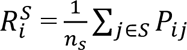, where *n_s_* is the number of nodes in the network *S*. *P_ij_* represents the number of times that nodes *i* and *j* are assigned to the same community. Dynamic integration was defined as the probability that a region is assigned to the same network community as regions from other networks across layers and repetitions. For node *i* in the community network *S*, the dynamic recruitment can be computing as follows: 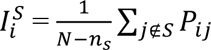, where *N* is the total number of brain regions. For each participant, both recruitment and integration coefficients obtained for each brain regions were averaged by networks to quantify brain networks’ dynamic recruitment and integration over time (Figure 3C, bottom). Although the dynamic recruitment and integration coefficients were computed based on all 68 regions from the six predefined networks, we focused the analyses and visualizations on the three core cognitive networks (DMN, SN, and CEN) due to their critical roles in cognitive functioning and aging-related changes. This selection aligns with prior literature highlighting these networks as central to higher-order cognitive processes and their sensitivity to aging and neurodegeneration (Menon, 2011, La Corte et al., 2016). Module allegiance matrix, enables to visualize brain networks’ dynamics (Mattar et al. 2015), as the diagonal of the matrix represents the dynamic recruitment and the off-diagonal represents the dynamic integration (Bassett et al. 2015) (Figure 3C, top and bottom).

### 5.9. Statistical analyses

To statistically assess the differences between the DMN, SN and CEN networks recruitment and integration dynamic measures over time in theta and gamma bands, 3 (*networks*: DMN, CEN, SN) x 2 (*dynamic measures*: integration, recruitment) x 2 (*frequency bands*: theta, gamma) repeated-measures ANOVAs with subjects as a random factor were applied using JASP (https://jasp-stats.org/; version 0.18.1). The Greenhouse-Geisser epsilon correction was applied to the factor *networks* where Mauchly’s test indicated a violation of the sphericity assumption. Original degrees of freedom and corrected *p*-values are reported. Finally, regressions aimed at determining the association of dynamic recruitment and integration measures of each network in theta and gamma frequency band, with cognitive performances scores at baseline, cognitive decline scores (the three domain-specific composite scores and the recognition memory score of the MST) and β-amyloid rates and tau/neuroinflammation burden rates (the latter only available for 64 participants). Results were FDR corrected for multiple comparisons (Benjamini & Hochberg, 1995), with a significance threshold of q < 0.05 applied separately to each set of comparisons (2 comparisons for ANOVA, and 3 to 4 comparisons for each regression analysis set). Participants’ age, sex and mean gray matter volume and individuals’ wake-up/DLMO phase value were included as covariates in the analyses.

